# Exploring the limitations of event-related potential measures in moving subjects. Case studies of four different technical modifications in ergometer rowing

**DOI:** 10.1101/578534

**Authors:** Holger Hill

**Author notes:** Correspondence: Holger Hill, Karlsruhe Institute of Technology, Institute of Sports and Sports Science, Mental mHealth Lab, Hertzstr. 16, Geb. 06.31 (LS Psychologie), D-76187 Karlsruhe, Germany.

## Abstract

Measuring brain activity outside the laboratory is of great importance for investigating human behavior under naturalistic conditions (e.g., in cognition and movement research, application of brain-computer interfaces). To measure neuronal activity in moving subjects, only modified NIRS and EEG systems are applicable. Because conventional EEG systems are too sensitive to movement artifacts, artifact sources should be eliminated beforehand to improve signal quality. Four different approaches for EEG/ERP measures with moving subjects were tested in case studies: (i) a purpose-built head-mounted preamplifier, (ii) a laboratory system with active electrodes, (iii)+(iv) a wireless headset combined with (iii) passive or (iv) active electrodes. A standard visual oddball task was applied during rest (without movement) and during ergometer rowing. All 14 measures revealed very similiar (within subjects) visual evoked potentials for rowing and rest. The small intraindividual differences between rowing and rest, in comparison to the typically larger interindividual differences in the ERP waveforms revealed that ERPs can be measured reliably even in an athletic movement like rowing. The expected modulation of the motor-related activity by force output, on the other hand, was largely affected by movement artifacts. Therefore, for a successful application of ERP measures in movement research, further developments to differentiate between movement-related neuronal activity and movement-related artifacts are required. However, it cannot be excluded that activities with small magnitudes related to motor learning and motor control cannot be detected because they are superimposed by the very large motor potential which increases with force output.

## 1. Introduction

The investigation of brain functions with noninvasive methods like functional magnetic resonance imaging (fMRI), magnetencephalography (MEG), electroencephalography (EEG), near-infrared spectroscopy (NIRS), and transcranial magnetic stimulation (TMS) is typically limited to laboratory settings. This is because the systems are very large and cannot be moved and head movements must be avoided (fMRI, MEG) or head and body movements generate large movement artifacts. Within the last decade there has been a rapidly increasing interest in investigating brain functions in ecological settings (e.g., cognitive or neuropsychological processes in interaction with a natural environment or in a social context; and movement analysis and motor learning) which require portable systems (e.g., Ruffini et al., 2007; Chi et al., 2012; Gramann et al., 2011, 2014; Kranczioch et al., 2014; Stopczynski et al., 2014). Especially in the field of movement research/motor learning laboratory settings are strongly limiting because only simple finger or hand or arm movements can be investigated and it is questioned whether the results of these studies can be transferred to complex movements (e.g. Dum and Strick, 2002; Hazeltine and Ivry, 2002; Wulf and Shea, 2002). In principle, NIRS and EEG are suited for measures in moving subjects because the sensors are small and fixed to the head (rather than the head being fixed to the sensor) and the necessary electronics and recording devices can be built small enough. NIRS, like fMRI, measures the hemodynamic response related to specific brain processes with a low temporal resolution. Spatially it is restricted to cortical layers close to the skull. Atsumori et al. (2010) used a modified NIRS system successfully in a cognitive task when subjects were walking around. Piper et al. (2014) developed a miniaturised wearable functional NIRS system and tested it when subjects performed a left hand gripping task (i) sitting still on a bicycle, (ii) pedalling indoor on a stationary training bicycle, and (iii) during outdoor bicycle riding. The event-related data showed that the task was performed successfully and comparably in all three conditions, whereas data loss was highest in real cycling (about 35%), but much lower in indoor cycling (7.5%) and under rest (5%).

Event-related potential (ERP) measures, which are the focus of this paper, have the well-known limited spatial resolution but a high temporal resolution which is essential to analyze complex movements: for example the time course of feedback and feedforward processing in visuomotor learning (e.g. Hill, 2009, 2014). Conventional EEG systems are very sensitive to mechanical (cable and electrode movements) and physiological (electromyogram (EMG) of head and neck muscles, and sweating) movement artifacts (Zschocke, 2002). If cognitive processes in a moving subject rather than the movement itself are the focus of interest, data preprocessing algorithms informed by the behavioral movement data can be used to clean the EEG data from movement-related (neuronal and artifactual) activity (Gwin et al., 2010; Stone et al., 2018). If the motor-related activity is of interest, advanced data preprocessing algorithms such as independent component analysis (ICA; Makeig et al., 1996) can be used to correct such artifacts. However, a study measuring ERPs during walking and running on a treadmill showed that data loss was very high (on average 130 of 248 EEG channel signals remained), even using a system with active electrodes which are considerably less prone to artifacts than conventional passive electrodes (Gramann et al., 2010; Gwin et al., 2011). Furthermore, the usability of conventional EEG systems like the one used in these studies is limited for measures with moving subjects. In movement tasks with only marginal head movements, like cycling on an ergometer, these laboratory systems combined with ICA-based artifact correction can be applied succesful for EEG measures (Enders and Nigg, 2016). E.g. Enders et al. (2016) found in a high-intensive cycling exercise an increase in spectral power when the athletes were fatigued. If spectral changes of higher EEG frequencies (alpha to gamma) are in the focus of interest, like in the Enders et al. study, artifacts directly coupled to movement execution are outside of this frequency range because movement frequencies are considerably lower. However harmonics of these movement frequencies may occur which have to be considered. With fully moving subjects, in contrast, artifacts are more difficult to handle. In a cocktail party study (including eating, drinking, chatting, etc.) with ten subjects wearing self-made (noncommercial) wireless EEG headsets, about 40% of the data was lost due to artifacts in contrast to 4% in two laboratory studies (Gevins et al., 2012). Despite this high data loss, these studies revealed valuable ERP (Gramann et al., 2010; Gwin et al., 2011) or spectral EEG (Gevins et al., 2012) results. However, especially in movement research the method of choice is to avoid the generation of mechanical artifacts beforehand by technical modifications. This approach was used in the four pilot studies reported in this paper that tested the suitability of different technical solutions to measure ERPs during ergometer rowing (Fig. 1). Especially for analysing motor-related brain activity with ERPs, rowing is well suited. It is a cyclic movement with a high number of repetitions and the degrees of freedom of the movement are limited by the biomechanical constraints of the equipment (boat and scull/oar, or ergometer). Furthermore, in contrast to e.g. cycling or cayaking, the rowing movement is composed of different (more or less) distinct movement elements and movement frequency is lower (20-40 strokes/minute vs. 60-120 revolutions/minute in cycling). Finally, the biomechanical data (dynamics and kinematics of the movement, and boat and scull/oar movement), which are partly necessary for ERP analysis, can be measured meanwhile with relative ease.

**Fig. 1.**
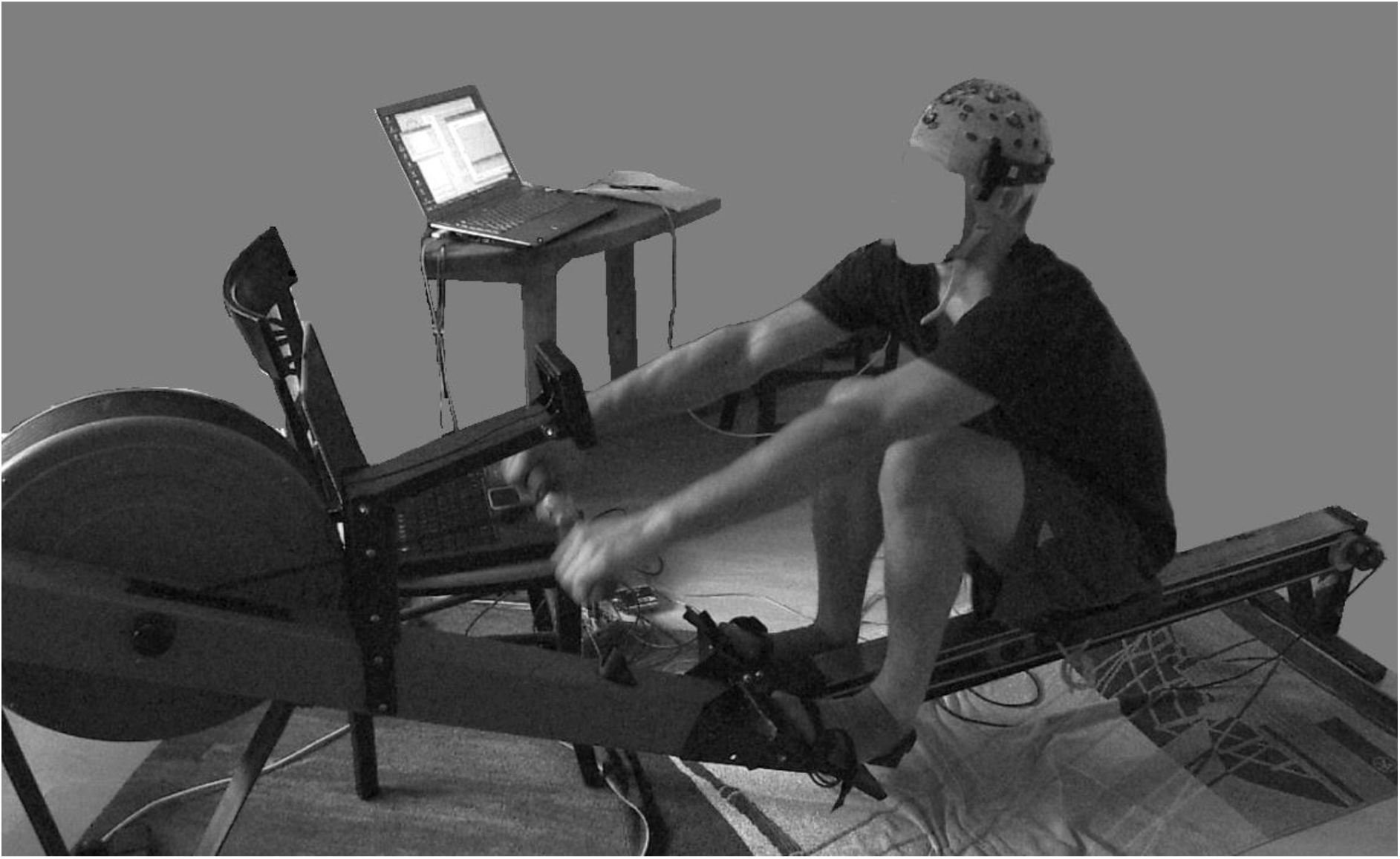
EEG recording during ergometer rowing in pilot Study 3. The Concept II (Model C) indoor rower was used for all three pilot studies. The Concept II models are the most frequently used ergometers for training in competitive rowing and performance diagnostics. In the drive phase of the rowing cycle, the rower’s pull accelerates an air resistance braked flywheel in the round cage. In the recovery phase the rower moves in opposite direction on the sliding seat, preparing for the next pull. A monitor displays stroke rate, time, and distance rowed, power per stroke, mean power, calories burned. On the chair at the right side in front of the rower is the laptop for stimulus presentation. The laptop on the table recorded the behavioral and EEG data (photo use with participant’s permission).

The most critical question before pilot Study 1 (see below) was, if movement artifacts distort the EEG data completely or if there are systematic artifacts which can be controlled and either (partly) excluded or corrected in offline analysis. To test this question, a reliability check was made. A standard visual oddball task was applied in a rest condition (without movement) and during ergometer rowing. Similar – not movement related – ERP activities during rest and during rowing would show that ERPs can be measured reliably in a moving subject. The second methodological question – concerning the analysis of motor behavior with ERPs – is: can movement-related artifacts be identified and separated from motor-related neuronal activity? As an indicator for a reliable measure of motor-related activity, at least a motor potential (MP), a negative activity related to force output, should be expected. Siemionow et al. (2000) showed in a study which used isometric elbow flexions that the amplitude of the MP (labelled motor-related cortical potential (MRCP) in this study) correlates very highly with EMG activity (r > 0.8) and the generated muscle force (r = 0.95). Furthermore, Dai et al. (2001) found in a fMRI study a high correlation between isometric force (using a hand-grip dynamometer) and activity in the primary motor cortex. The MP/MRCP must be generated by different generators because it can already be evoked when a muscle activation is only imagined. This should originate in the supplementary motor area (SMA; Ranganathan et al., 2004). A second part of the MP is thought to be related to the control of muscle activation by the primary motor cortex.

Objectives: The purpose of these four pilot studies was (i) to test if ERPs can in principle be measured during an athletic movement pattern like rowing, (ii) to explore the limitations due to movement artifacts, and (iii) to test if movement artifacts can be reduced. Methodologically, the impact of physical and physiological artifact sources was investigated by comparing ERP waveforms measured in rest and during movement. For the existing data material this physical approach is the method of choice compared to a statistical analysis of parametrized abstract data derived from the original data. Study outlines: In pilot Study 1 (conducted in 2005) a small, purpose-built 20-channel system was used with a preamplifier connected to the head and electrodes with shortened and fixed cables mounted to a standard electrode cap. Because this approach was only partly successful the second approach (Study 2) was performed. This experiment involved the use of a to that time (2008) in our lab available system with active electrodes with built-in preamplifiers, and the amplifier was worn in a backpack. In pilot Study 3 (2013), as the newest improvement in EEG technology, a small head-mounted EEG system with wireless data transmission was combined with the electrodes and cap used in Study 1. In Study 4 (2018), finally, the headset used in Study 3 was combined with active electrodes and rowing force and movement speed was systematically varied to investigate the influence of these factors on data quality.

## 2. Pilot Study 1

### 2.1. Materials and Methods

A preliminary test using a standard EEG system (NeuroScan SynAmps) by wearing the 32-channel preamplifier headbox in a backpack revealed no satisfying results. Therefore, to reduce the generation of artifacts due to cable and electrode movements, a small purpose-built (according to Simon, 1977) battery-powered 20-channel system (based on positive previous experiences with a similar three-channel system) was used with an occipitally mounted preamplifier (differential amplifier, gain = 30; hardware filters: 0.27 Hz passive RC highpass; 30 Hz 2nd-order Bessel lowpass). After a second amplifier stage (total gain = 3600) the signals were digitised using a BEST system (Dr. Grossegger & Drbal, Korneuburg, Austria; sample rate 256 Hz, resolution 0.3 μV/bit). An electrode cap (EasyCap, www.easycap.de) and 22 gold electrodes (Grass) with shortened cables were used. Electrode caps with electrodes fixed to the cap have the disadvantage that the contact of some electrodes with the skin can be poor, depending on head shape (that is, if there are dents in the skull the distance between cap and skin can be too large). Therefore the standard adaptors inserted in the cap were removed, the adaptor holes were enlarged, and the electrodes were fixed using a highly viscous conductive paste (Nihon-Kohden Elefix). EEG was recorded from 20 sites (midline: FPz, Fz, FCz, CPz, Pz, and Iz; Left/right: FC1/2, FC3/4, FC5/6, C3/4, CP1/2, CP3/4, left and right mastoid) covering mainly the sensorimotor area, and Iz to assess visual evoked potentials (VEPs). Data were recorded using Cz as reference and rereferenced offline (see results for details). A visual oddball task was used with 180 black-white fullscreen checkerboards (frequent), 60 red-white checkerboards (deviant), and 60 gray crosses (target) which had to be counted. Stimulus duration was 100 ms, the interstimulus interval varied randomly between 800 and 1000 ms. The oddball task is a standard paradigm and generates robust ERP components: VEPs at occipital sites which are generated in the visual cortex when visual stimuli are presented. The P300 at centroparietal sites (around CPz, Pz) when attention is shifted to a target stimulus. Stimuli were presented asynchronous to the temporal pattern of the rowing movement on a computer screen besides the ergometer using the Presentation software (Neurobehavioral Systems, www.neurobs.com), allowing to monitor the display without head movements, although eye movement artifacts may be generated. Alternatively, an acoustical stimulation could be used. However, because the auditory cortex is closer to the sensorimotor cortex and to the mastoids than the visual cortex, this will probably lead to extended signal overlay of motor-related and auditory-evoked potentials. The EEG was analyzed using the Vision Analyzer 1.0 software (BrainProducts, Gilching, Germany). EEG data were digitally filtered with a 16 Hz/24 dB Butterworth zero phase lowpass, segmented into epochs of −200 ms to 800 ms around stimulus onset (oddball task), and baseline corrected (−200 to 0 ms). After applying a semiautomatic procedure for artifact detection (amplitude criterion ±100 μV, gradient 25 μV/sample) the complete datasets were inspected visually for further artifacts surviving the automatic rejection. Segments with blink or eye movement artifacts (detectable at frontal sites) were excluded completely. Traces of single channels containing other clearly visible large artifacts were removed. If there were more than five contaminated traces, the whole segment was removed. ERPs were computed for each stimulus category of the oddball task. Motor-related ERPs were computed triggered by the force onset at the beginning of the rowing stroke. Rowing force was measured with a modified ergometer handle using strain gauges (according to the measure of oar forces in rowing; Hill, 2002) and movement of the sliding seat was measured with a potentiometer. Biomechanical data were recorded with a second computer, synchronised using triggers of the Presentation software, and added to the EEG data file offline.

Before performing this pilot study, a statement of the ethics committee of the German Society of Psychology was obtained for a grant proposal. This stated no ethical concerns about this type of investigations. Written informed consent was obtained (respectively from a parent when under 18). Three male subjects (aged 47, 13, and 10 years, and referred to here as: H, J, and M respectively) performed (i) the oddball task sitting still on the ergometer, (ii) rowing without stimulation, and (iii) the oddball task during rowing. Subjects were instructed to row in a recreational mode (e.g., 130 W for subject H) during the oddball task and to keep rowing power and stroke rate (20/min) constant, using the performance monitor of the ergometer for control. The rest condition of the oddball task was always performed first to avoid sweating artifacts.

### 2.2 Results

Fig. 2 displays ERPs of all three subjects for the oddball task. Especially the VEPs were very similar between rowing and rest, and intraindividual differences were much smaller than the interindividual differences. This result demonstrates that standard ERPs not time-locked to the rowing movement can be measured during rowing, despite the fact that subjects M and J had no or only marginal rowing experience. In contrast, the motor-related ERPs were very noisy and showed large artifacts with large inter- and intraindividual (implemented by varying force output and stroke rate) differences and were therefore not interpretable. One observable source of large artifacts was due to movements of the cable connecting the head-mounted preamplifier with the second amplifier unit, which led to movements of the preamplifier and electrode cables.

**Fig. 2.**
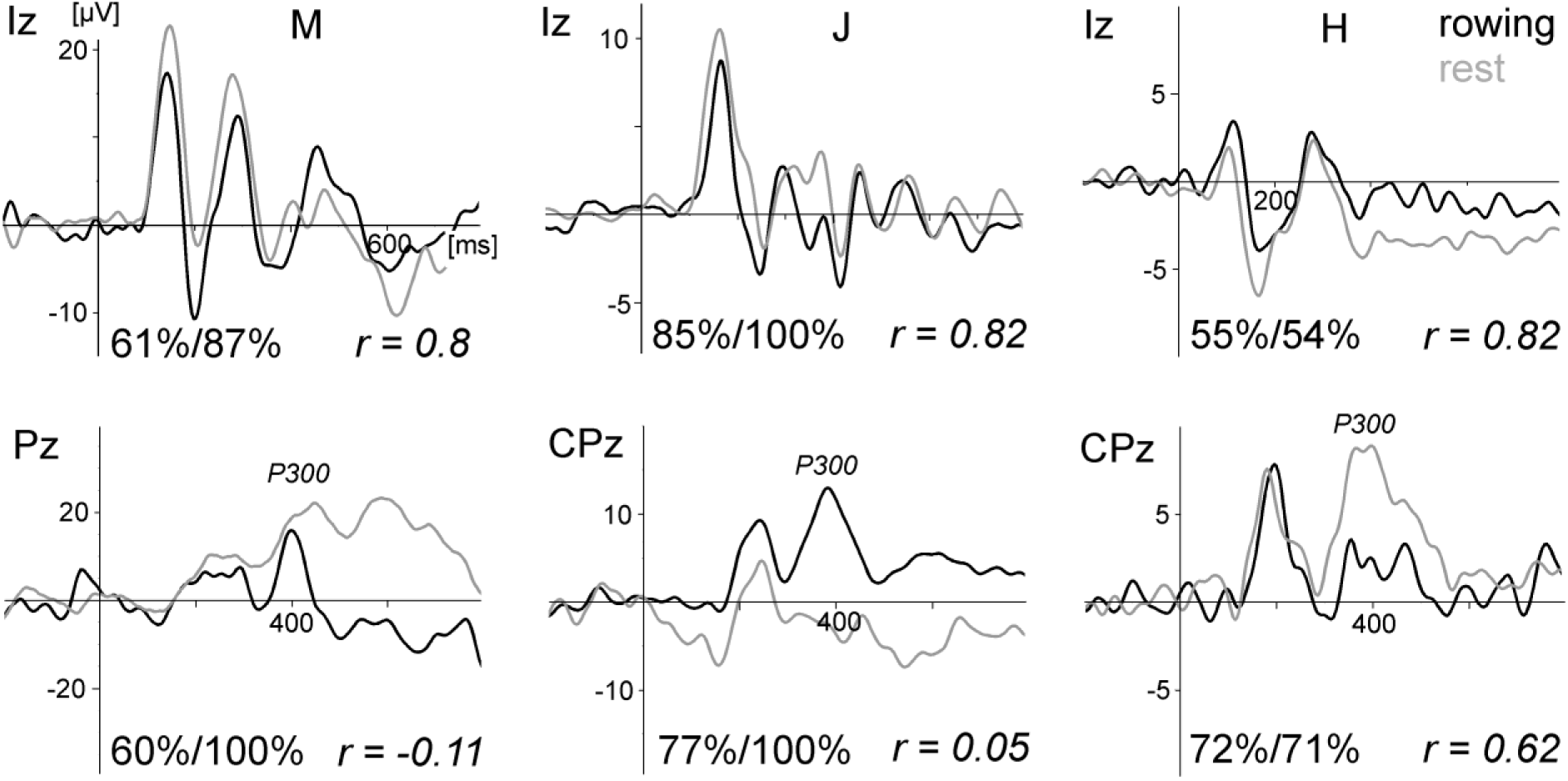
ERPs of the visual oddball task comparing rowing and rest for all three subjects (M, J, and H) of pilot Study 1. Upper graphs: VEPs evoked by the frequent checkerboard stimulus at electrode site Iz (referenced to Cz to obtain a larger and cleaner signal because VEP activity and EMG artifacts are lower at Cz compared to the mastoids). Below: the P300 waveform at Pz/CPz (linked mastoid reference for M and H, and right mastoid for J due to the lost left mastoid channel). The vertical line marks the stimulus onset. The numbers at the bottom of each graph present the percentage of artifact-free trials included in the averages for rowing and rest of the chosen electrode site and the correlation coefficients (Pearson) of the two waveforms. The high dropout rate for subject H was mainly due to eyeblink artifacts. For subject M three channels (FCz, CPz, and CP4) were lost completely due to artifacts during rowing: for subject J the left mastoid channel was lost. Furthermore, a P300 is missing during rest for J because he did not count the target because of an imprecisely given instruction. For M and H the P300 was larger during rest. The ERPs of the deviant stimulus condition are not displayed because they showed results similar to those of the frequent condition.

## 3. Pilot Study 2

### 3.1. Materials and Methods

The second approach used a system with active electrodes at the Department of Psychology, University of Frankfurt. This system suppresses artifacts due to cable movements, however electrode movements cannot be avoided completely. During rowing and rest the amplifiers were worn in a backpack and connected via fibre-optic cables to a PC. EEG was recorded continuously with BrainAmp DC amplifiers (BrainProducts, Gilching, Germany; sample rate 250 Hz, resolution 0.1 μV/bit, input impedance 10 MOhm) using an equidistant EasyCap (EasyCap GmbH, Herrsching-Breitbrunn, Germany, www.easycap.de) with 62 sintered Ag/AgCl electrodes and built-in preamplifiers (BrainProducts ActiCap System). Eye blinks and movements were monitored with supra- and infra-orbital electrodes and with electrodes on the external canthi. The vertex electrode was used as the reference. To avoid injuries due to skin abrasion, electrode impedances were kept at 20 kOhm which is more than sufficient from electrical engineering principles (Ferree et al., 2001; see Picton et al., 2000). The EEG was analyzed like in Study 1 with slightly different filter settings (0.5 to 20 Hz bandpass). The averages were rereferenced (average reference transformation; Bertrand et al., 1985), and the reconstructed vertex reference was added to the data, resulting in 61 EEG channels. The ergometer and its measuring equipment and the experimental procedure were the same as in Study 1. The task was performed by two female students (F and K, aged 24 and 26), without rowing experience and an experienced male rower (H, aged 50). Written informed consent was obtained. Subjects rowed in a recreational mode during the oddball task with a stroke rate of 20/min.

### 3.2. Results

Fig. 3 displays ERPs of all three subjects for the oddball task. As in Study 1, the VEPs were very similar between rowing and rest, the P300 was smaller during rowing, and the intraindividual differences were much smaller than the interindividual differences.

**Fig. 3.**
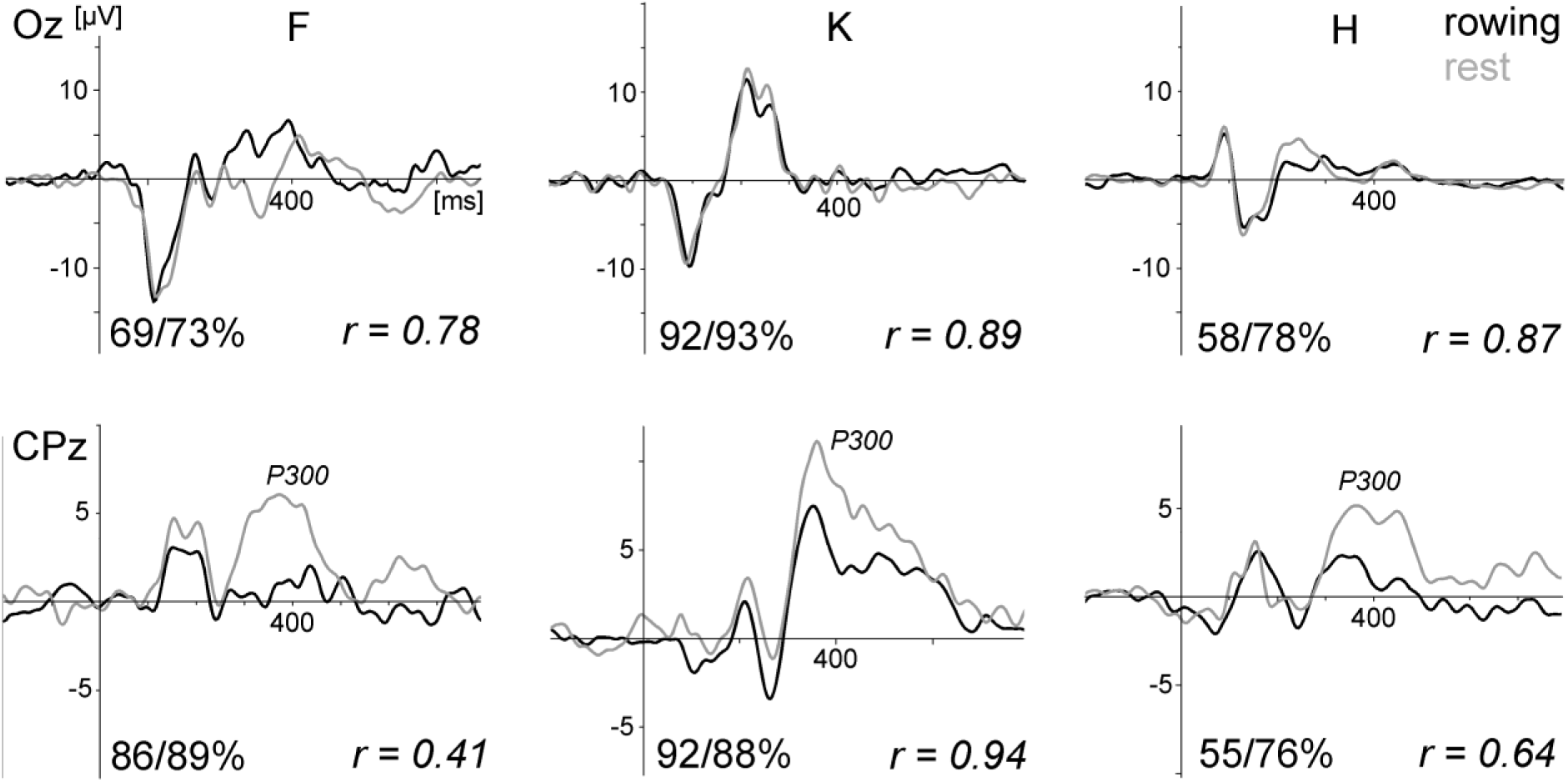
ERPs of the visual oddball task comparing rowing and rest for all three subjects (F, K, and H) of pilot Study 2. Upper graphs: VEPs evoked by the frequent checkerboard stimulus at electrode site Oz (average referenced). Below: the P300 waveform at CPz, which was larger during rest. The vertical line marks the stimulus onset. The numbers at the bottom of each graph present the percentage of artifact-free trials included in the averages for rowing and rest of the chosen electrode site and the correlation coefficients. The high dropout rate for subject H was due to eyeblink/eyemovement artifacts. After an ICA-based correction, about 98% of the trials could be kept, resulting in very similar ERPs. For subject F the data loss at occipital sites was due to neck muscle activity. Deviant stimulus condition is not shown.

Motor-related activity, on the other hand, was not dominated by the expected motor potential in all conditions. Applying an ICA-based artifact rejection procedure improved the data quality somewhat but insufficiently, probably because movement artifacts are not stable enough over time, which is a prerequisite for computing the ICA components. Performing only the arm pull revealed for subject H a bilateral negativity; however, for subjects F and K it revealed no interpretable results. For normal rowing a negative activation at central sites, indicating a motor potential, appeared, but at peripheral electrode sites large activities occurred which were only partly corrected using ICA. One possible source for artifacts in this study were electrode movements due to cable drag because cables were not fixed to the cap.

In summary, this pilot study revealed results similar to those of Study 1. Standard ERPs can be measured reliably during rowing, however motor-related activity is largely distorted by remaining artifact sources.

## 4. Pilot Study 3

### 4.1. Materials and Methods

Disadvantages of the two systems used in the previous pilot studies were the amount of required equipment and the data transmission via cable connections. Therefore, a study of Debener et al. (2012) was very promising. They used a new developed wireless system (Emotiv, www.emotiv.com). After making some technical improvements they measured ERPs in subjects walking around. The Emotiv system integrates the hardware in a small and lightweight headset, in combination with a wireless data transmission via an USB dongle to a laptop or even an Android smartphone (Stopczynski et al., 2014). Furthermore, this system integrates a two-axis gyroscope which measures head rotations. Electrodes and cables of the Emotiv system are fixed to stiff plastic arms which should be effective to reduce artifacts by cable and electrode movements. Acceptable limitations of the Emotiv system, at least for pilot studies, are the fixed and lower sample rate and resolution and the lower number of channels (14). In addition, the sensor locations cannot be changed and are not well suited to sensorimotor research. Therefore, as already practised by Debener et al. (2012), this system was modified by removing the original sensors and plastic arms and connecting the system to the electrode cap with the gold electrodes used in Study 1 (Fig. 4). Comparison studies revealed that the Emotiv system (equipped with the cheap original electrodes) performs less accurate than a medical device, however is able to record EEG data in a satisfying manner (Duvinage et al., 2013; Badcock et al., 2013). Comparing a modified (according to Debener et al., 2012) Emotiv System with a commercial SynAmps System revealed only small differences, with a marginal worse performance of the modified Emotiv System (Barham et al., 2017).

**Fig. 4.**
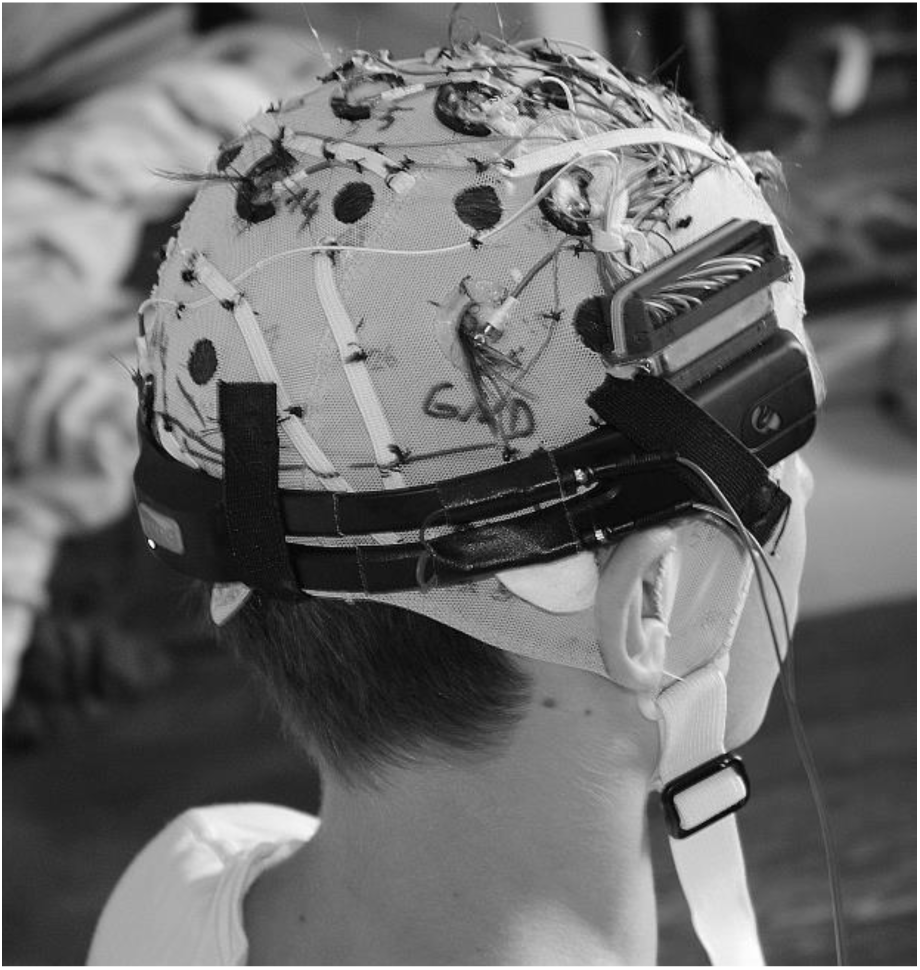
The modified Emotiv system combined with a standard electrode cap. A connector was fixed above the part of the headset containing the amplifier circuits. The two cables connected the headset with the photodiode used for synchronisation of stimulation and EEG recording at the start and end of each experimental run. During rowing the cables were removed.

A 14-channel EEG covering mainly the sensorimotor area (electrode sites AFz, Fz, FCz, Cz, CPz, Pz, FC1, FC2, CP1, CP2, C3, C4, O1 to assess VEPs, left mastoid, and reference right mastoid) was recorded using the Emotiv system (fixed settings: sample rate 128 Hz, resolution 14 bit/0.51 μV, bandpass filter 0.2 - 45 Hz, notch 50/60 Hz, digital 5th order Sinc filter, data transmission 2.4 GHz band, and operation distance measured outside about 10 m). Signal quality was controlled with the Emotiv Testbench recording software. The ergometer, its measuring equipment, and the oddball task (except a reduction of the number of standard stimuli from 180 to 120) were the same as in Study 1. The synchronisation of the EEG data with the biomechanical data and the visual stimulation was somewhat difficult. Although the Emotiv system can read trigger signals from a serial port, the laptops used did not have serial ports. An interface with serial-to-USB adaptors is not accurate enough in timing. Therefore, the Presentation scenarios were modified for synchronisation. A photodiode was fixed to the Presentation laptop and activated by stimuli at the beginning and the end of each experimental run. The triggers sent via the parallel port were recorded with an USB analog-digital device (RedLab 1208 LS, Meilhaus, Puchheim/Germany) together with the biomechanical data (force of the rowing stroke and movement of the sliding seat). The signal of the photodiode (which was much larger than the EEG) was recorded with one EEG channel, and the cables were removed during the experimental runs. The EEG, biomechanical data, and triggers of the oddball stimulation were synchronised offline using purpose-written software. A separate channel for received data packets implemented in the Emotiv system provides the ability to control for lost data. This infomation was used to correct the synchronisation (a small fraction of samples was lost in 4 of 25 datasets). The EEG was analyzed in a way similar to that used in Study 1 (using only a 20 Hz lowpass filter). Data of the rowing condition of two subjects had a larger number of trials contaminated with eye blinks. These were corrected successfully using ICA without affecting other activities (despite the low spatial resolution relying on only 14 channels can be critical to apply the ICA procedure).

Four male subjects (aged 55, 21, 18, and 14 years, and referred to here as H, J, M, D), with ergometer rowing skills allowing a high performance were instructed to perform six experimental conditions as follows: (i) oddball-rest; (ii) rowing with arm pull only (power 50 W, stroke rate (SR) 30/min); (iii) rowing 100 W/SR 20/min; (iv) oddball-rowing 100 W/SR 20/min; (v) rowing 180-200 W/SR 20/min; (vi) rowing 180-200 W/SR 26/min. Power and stroke rates could be controlled with the monitor of the ergometer and were close to the instruction. The rowing power ranged from recreational to long distance endurance rowing. For subject D the planned power thresholds were reduced to 60%. Written informed consent was obtained (respectively from a parent when under 18).

### 4.2. Results

Fig. 5 displays ERPs of all four subjects for the oddball task. As in studies 1 and 2, the VEPs were very similar between rowing and rest, the P300 was smaller during rowing, and the intraindividual differences were much smaller than the interindividual differences.

**Fig. 5.**
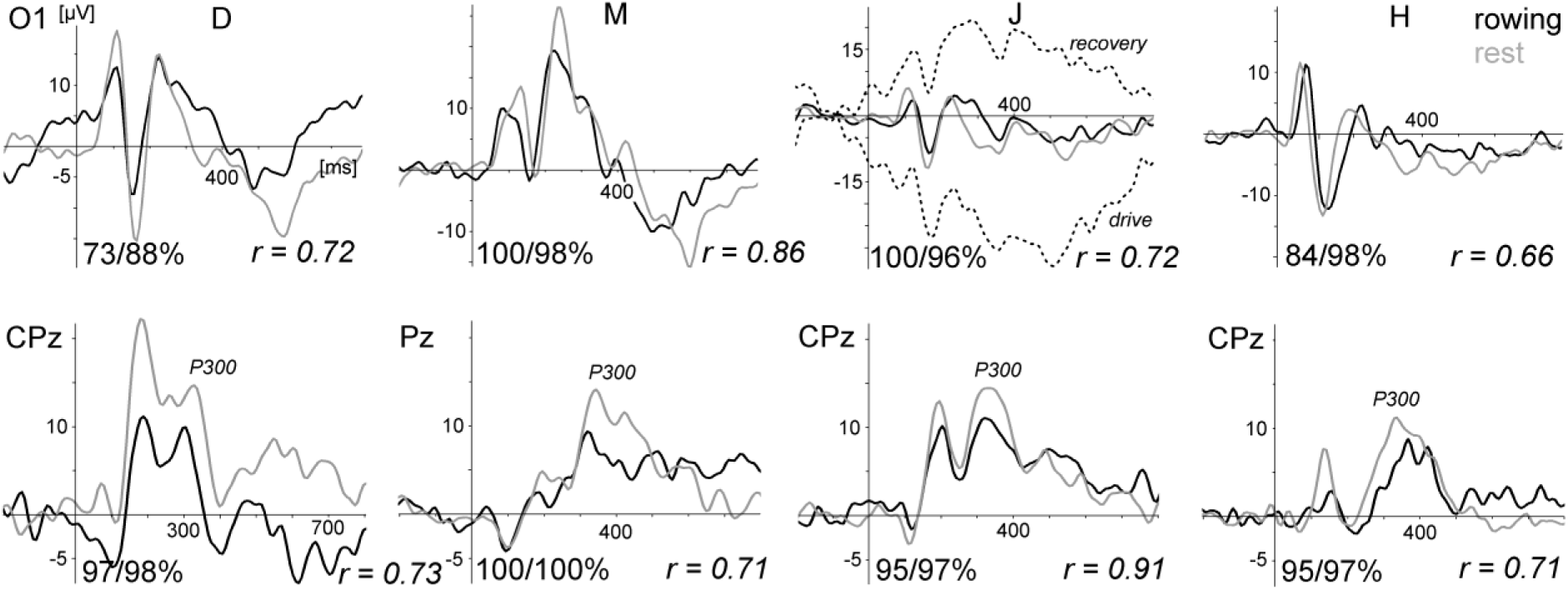
ERPs of the visual oddball task comparing rowing and rest for all four subjects (D, M, J, and H) of pilot Study 3. Upper graphs: VEPs evoked by the frequent checkerboard stimulus at electrode site O1 (referenced to Cz). Below: the P300 waveform at CPz/Pz (referenced to linked mastoids) which was larger during rest. The vertical line marks the stimulus onset. The numbers at the bottom of each graph present the percentage of artifact-free trials included in the averages for rowing and rest of the chosen electrode site and the correlation coefficients. Deviant stimulus condition is not shown. For the frequent stimulus of subject J, the ERPs of the rowing condition, separated into drive (with force) and recovery phases of the rowing movement (without excluding artifactual trials), are presented additionally. In this recording the VEP at electrode site O1 was overlaid by large movement-related drift artifacts, which cancelled each other out when all trials (drive and recovery) are included in the average waveform.

The motor-related activity showed more or less large artifacts at several sites, as well as the expected modulation by rowing force at some other sites – that is a larger negative activity with increasing force during the rowing stroke – with some differences between subjects (Fig. 6). As it can be difficult to differentiate between motor-related activity and movement-related artifacts at sensorimotor sites, movement-related artifacts can clearly be identified at sites outside the sensorimotor region. Fig. 5 displays such an example for subject J at electrode site O1. The VEP for the frequent stimulus of the oddball task was computed separately for the drive and recovery phases of the rowing movement (stimuli presented in the transition drive-recovery were excluded for this analysis). These waveforms were overlaid by large movement-related artifacts with reversed polarity which cancelled each other out in the average including all trials. These artifacts were independent of the chosen reference (Cz, right mastoid, and linked mastoids). That is, the overlay does not depend on physiological motor activity, which is much stronger at Cz than at the mastoids and O1; instead, it must be generated at electrode O1. Probably this was due to cable artifacts in an electromagnetically noisy environment. Furthermore, the electrodes O1, left mastoid, and AFz had the largest distance to the Emotiv connector and therefore the longest cables, which may have allowed small movements.

**Fig. 6.**
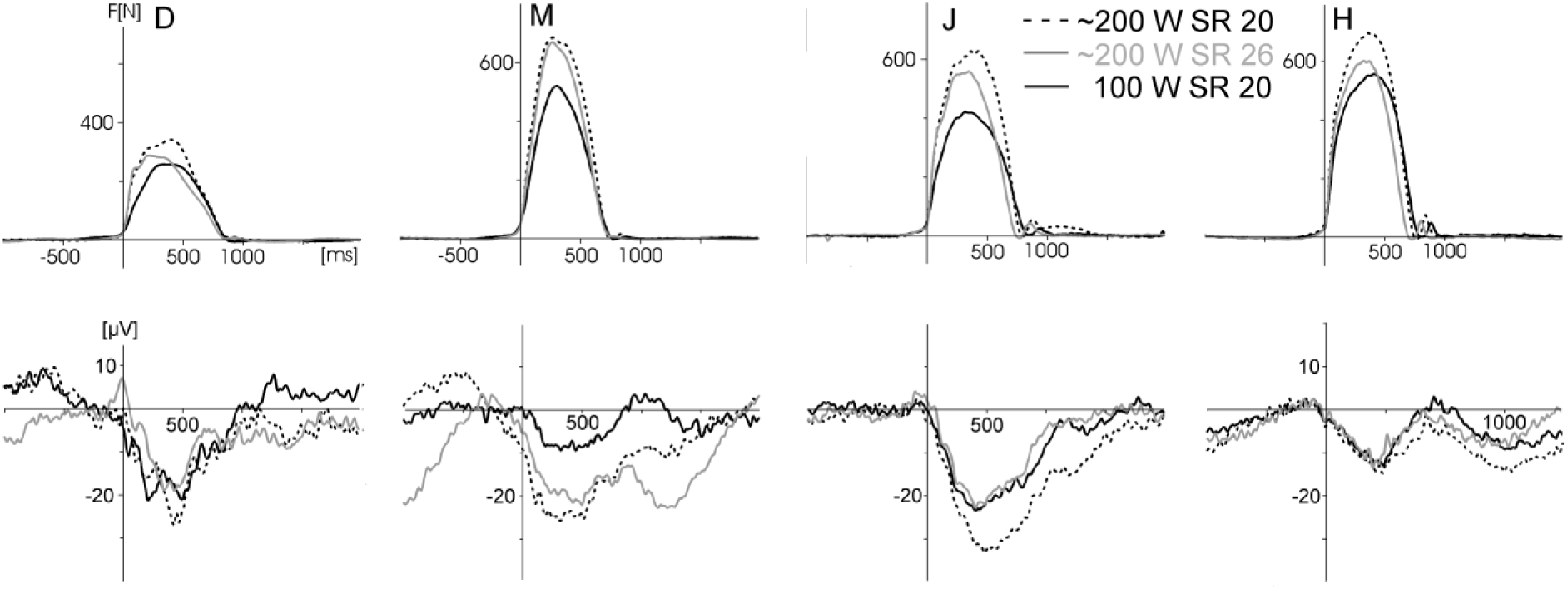
Force graphs (top row) and electrical activity (below) for the three different rowing conditions for all four subjects (D, M, J, and H) of pilot Study 3. Black solid: power 100 W, 20 strokes/min. Black dotted: 180-200 W, 20 strokes/min. Gray: 180-200 W, 26 strokes/min (lower power values for D). The electrical activity reflects the motor-related ERP (triggered by force onset, baseline −500 ms to 0) plus movement-related artifacts. Six bilateral sites (FC1, C3, CP1, FC2, C4, and CP2, referenced to the right mastoid) were averaged for this figure. The number of rowing cycles included in the different averages were between 54 and 88. Only a few EEG segments showing excessive and clearly identifiable artifacts (e.g., voltage jumps) were excluded.

To identify one possible source of movement artifacts, the gyroscope of the Emotiv system was used to test if artifacts are caused by head movements. Therefore, rapid repeated movements were performed (left turn, right turn, and nodding) revealing very large artifacts, especially at lateral sites where the impact of the head movement was larger than at central sites. Using ICA these artefacts could be strongly attenuated (Fig. 7). A comparison of the gyroscope data for these head rotations and rowing showed only small head movements for nodding during rowing: that is, head rotations and the associated artifacts are not critical for rowing.

**Fig. 7.**
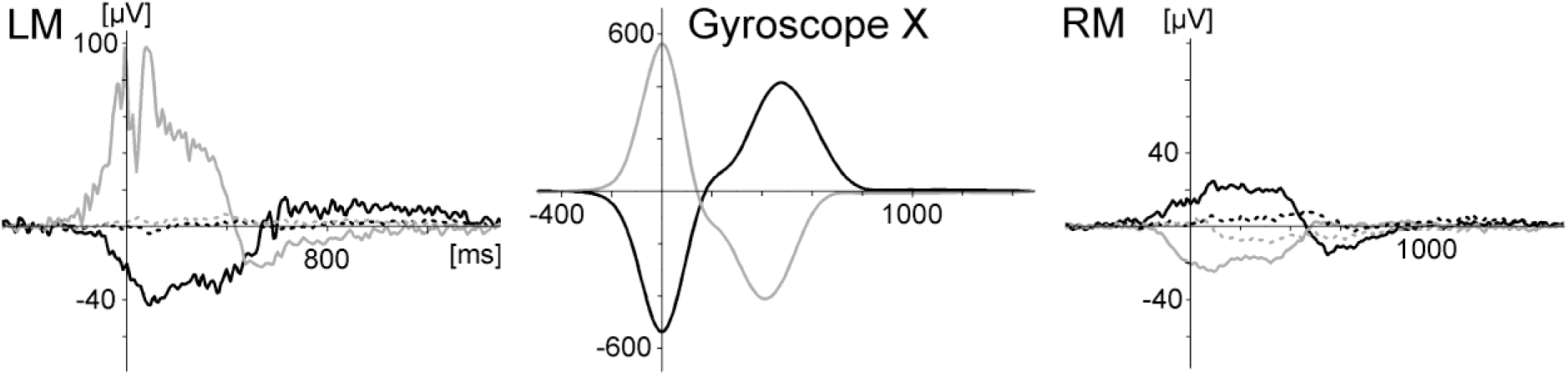
Movement artifacts generated by repeated rapid head rotations (measured with the gyroscope) at left mastoid (LM, where the artefact was largest) and right mastoid (RM, both sites referenced to Cz). Rotations to the right side: black lines; rotations to the left side: grey lines. The polarity was inverted between left and right electrode sites and between left and right head rotations. This artefact was present (with comparable amplitudes) at left sites C3 and O1, at right sites C4, FC2, CP2, and RM, and at O1, Fz, and AFz. Using ICA-based correction the artefact could be removed nearly completely (dotted lines). The gyroscope was not calibrated (arbitrary units).

In summary, this pilot Study 3 revealed results similar to those of Studies 1 and 2. Standard ERPs can be measured reliably during rowing, however motor-related activity is largely distorted by remaining artifact sources, whereas the use of, although modified, passive electrodes may be a limiting factor.Fig. 4. The modified Emotiv system combined with a standard electrode cap. A connector was fixed above the part of the headset containing the amplifier circuits. The two cables connected the headset with the photodiode used for synchronisation of stimulation and EEG recording at the start and end of each experimental run. During rowing the cables were removed.

## 5. Pilot Study 4

Under combining the technical advantages of Study 2 (active electrodes) and Study 3 (mobile headset), Study 4 followed two aims: (i) can movement artifacts be reduced when the passive electrodes of Study 3 are replaced by active electrodes, and can movement-related ERPs be measured in sufficient quality then? This would be the optimal result. (ii) If the first aim is not achieved: how is the signal quality of standard VEPs affected by movement dynamics (force output) and movement kinematics (movement speed). That is, is there a trade-off between signal quality and movement intensity in the measurement of ERPs in e.g. cognitive tasks in moving subjects.

### 5.1 Materials and Methods

The Emotiv system used in Study 3 was combined with eight active electrodes (EasyCap active) which were provided by EasyCap GmbH (www.easycap.de) together with electrode caps (EasyCap). In contrast to the older version of the Acticap electrodes used in Study 2 these electrodes are smaller and very flat, and should therefore be less susceptible to tilting movements generated by inertial forces. After pretests electrode cables were shortened, fixed to the cap and connected to the headset. Power for the electrodes was provided by a 9 V battery attached to the connector (Fig. 8). The electrodes were fixed with adaptors and a highly viscous conductive paste (Nihon-Kohden Elefix). The active electrodes were placed at positions Cz, FCz, C4', C3', O1, left and right mastoid (TP9, TP10), focusing the measure of motor-related activity and the VEP. The active reference electrode was placed at AFz for different reasons: (i) by generating eye blinks it could be controlled if all electrodes worked properly, (ii) sections with eye blink and vertical eye movement artifacts could be detected and removed from the EEG data, (iii) a mastoid reference is largely affected by EMG artifacts generated by neck and head muscles, (iv) the spatial distance to occipital sites is largest and allows to measure larger VEPs. A disposable baby-ECG electrode, which served as ground/DRL, was placed at the right anterior temple. The original passive ground electrode of the EasyCap-active system was used as a signal electrode and placed at Oz. Of course it is unusual to combine a passive electrode with an active reference electrode and it was not sure if it works because the amplifier would saturate if the impedances are very different. However this electrode always provided a clear EEG signal and no saturation effects were observed in any measure. Therefore the quality of the VEP measured with the active electrode at O1 could be directly compared with the VEP measured with the passive electrode at Oz. Signal quality was controlled with the Emotiv Testbench recording software. The recording parameters (A-D rate 128 Hz, bandpass filter 0.2 - 45 Hz) ergometer and its measuring equipment, and the oddball task were the same as in Study 3 except two modifications. The condition with the 60 deviant stimuli of the previous studies was removed because the VEPs did not differ from the standard checkerboard task. The number of trials of the latter was therefore increased from 120 to 180. Secondly, because from one subject no VEP signal was obtained, a second recording session was conducted at a later time (with a nearly identical rowing performance). To ensure that the visual stimulation was observed, the target stimulus (cross) was replaced by a lower number (27-33) of pictures of different airplanes. This was done because the oddball task was repeated six times instead of two times in the previous studies and therefore more salient stimuli were used. In this Study 4 the target condition of the oddball task was only used to control performance.

**Fig. 8.**
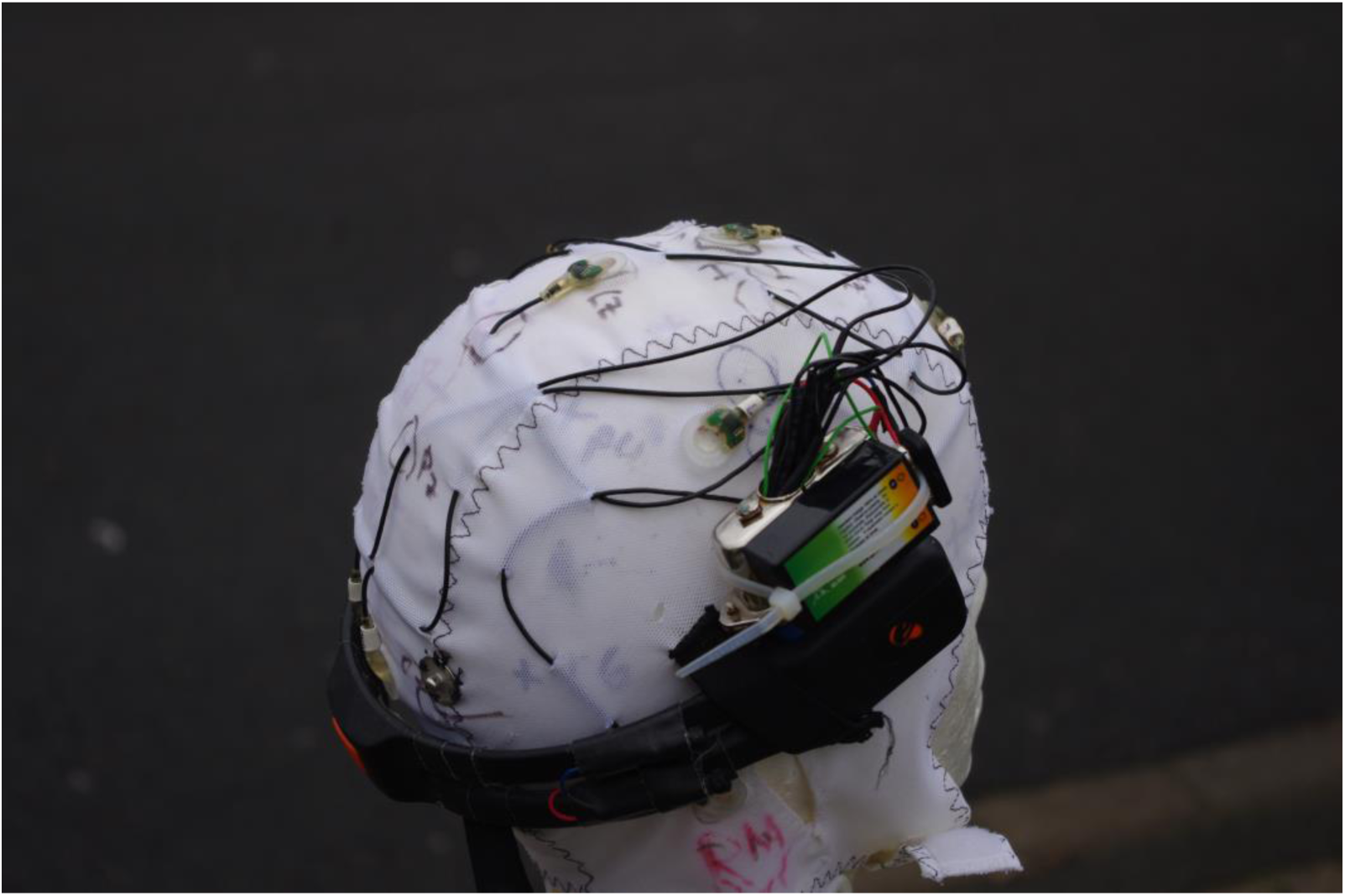
The modified Emotiv headset combined with the EasyCap active electrodes

**Fig. 9a.**
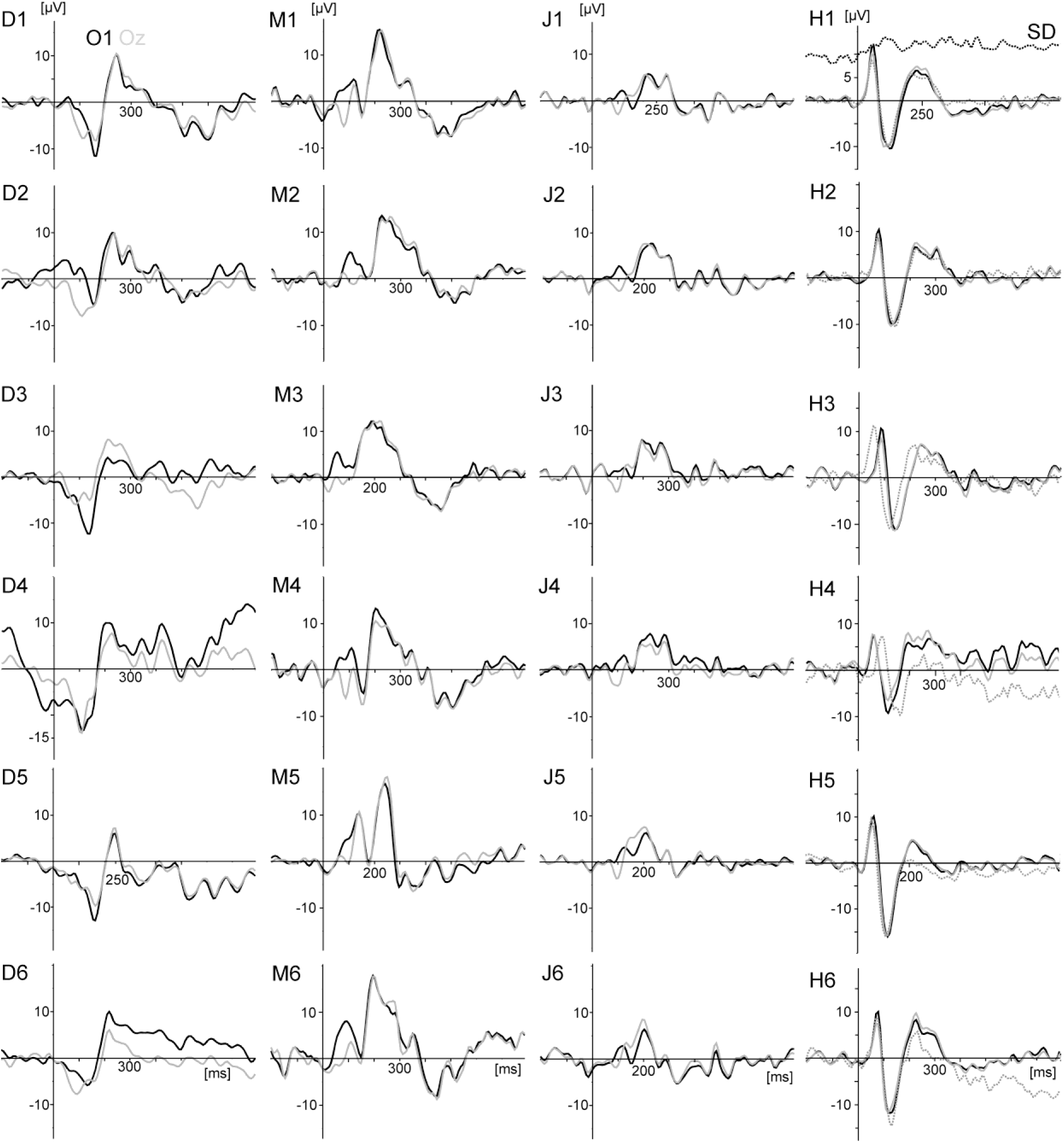
VEPs generated by the standard stimulation (checkerboard) for all four subjects (columns, D, M, J, H) and the six rowing conditions (rows). The VEP measured at active electrode O1 and passive electrode Oz are superimposed. In addition, for subject H (right column) the VEP at O1 measured in the repeated test with the 14-channel cap with passive electrodes used in Study 3 displayed (dotted grey line). In the upper right plot the standard deviation (time course) was included as an example. The six conditions were: 1: Force low (F_low), stroke rate low (SR_low). 2: F_low, SR_high. 3: F_high, SR_low. 4: F_high, SR_high. 5: Rest (without rowing). 6: Armpull only. Details are provided in the legend of Fig. 10

**Fig. 9b.**
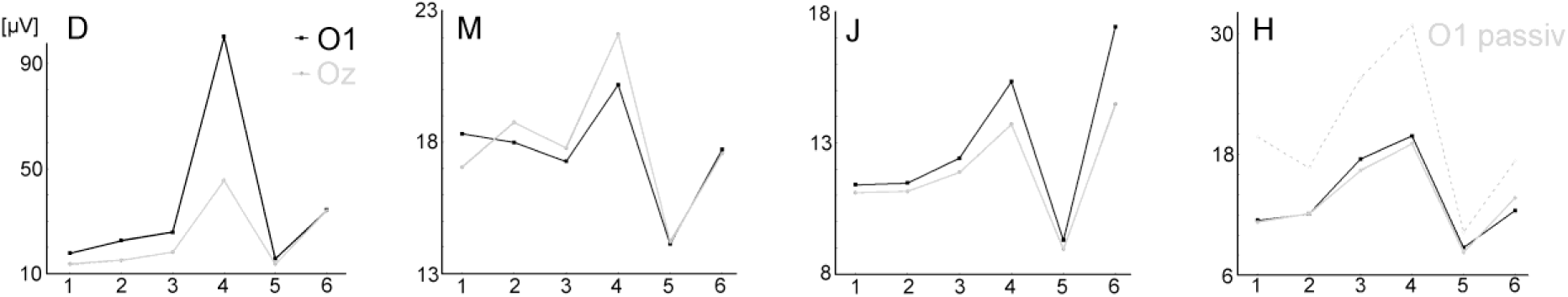
Standard Deviations (mean values) separately for conditions 1-6 and the four subjects.

The same four male subjects as in Study 3 (aged 19, 23, 26, and 59 years, height 180-190 cm, weight 80-84 kg) performed six experimental conditions of about four minutes duration comparable to Study 3: (i) rowing with lower force output and lower stroke rate; (ii) lower force output, higher stroke rate; (ii) higher force output, lower stroke rate; (iv) higher force output, higher stroke rate; (v) visual stimulation in rest (without rowing); (vi) rowing with arm pull only (details are provided in the legend of Fig. 10). Power and stroke rates could be controlled with the performance monitor of the ergometer and were close to the instruction. The rowing power ranged from recreational (condition i) to long distance (e.g. 10 km) racing (condition iv). The visual stimulation was applied in all six conditions. For comparison, one subject repeated all six conditions with the cap used in Study 3 with 14 passive electrodes.

**Fig. 10.**
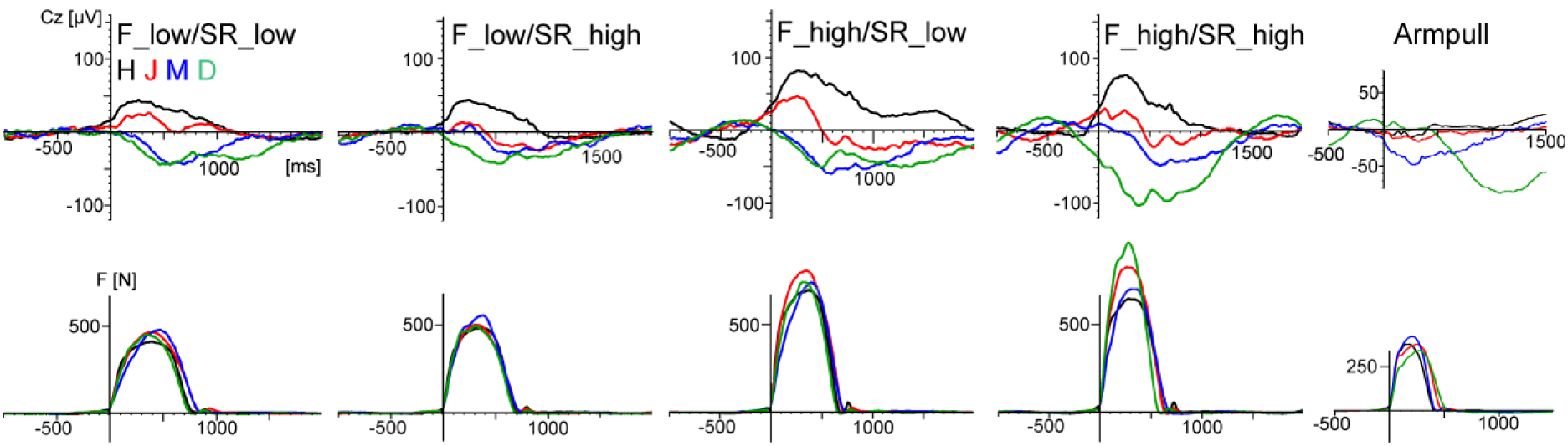
The upper row displays the force-onset triggered individual averages (subjects D, M, J, H) of the ERP at Cz (referenced to AFz) and at the bottom the averaged force graphs for the five rowing conditions (F force, SR stroke rate). ERPs at C3 and C4 were very similar to Cz and therefore omitted in this figure. The ERPs showed within-subject similarities with a higher activity in the conditions three and four with the higher force output, however large between-subject differences. The waveforms of subject D were largely affected by sweating artifacts in conditions four and six. Characteristics of rowing performance. Condition one: F_low, SR_low (individual mean values, range 19.4-21 min-1, 98-102 W). Condition two: F_low, SR_high (25.5-27.6), 121-163 W. Conditon three: F_high, SR_low (19.8-21.3), 159-192 W. Condition four: F_high, SR_high (26.1-28.6), 225-270 W. Condition six: armpull only (stroke rate 26.2-29.1), 50-57.5 W.

Data processing was performed as in Study 3, using Vision Analyzer 2.1 (20 Hz lowpass filtering, segment lengths 1 s for VEPs, 3 s movement related, 2 s condition 6). VEPs were analysed at electrode sites O1 and Oz (referenced to Cz like in Study 3). The number of included trials (total 180) was on average 138 (O1) and 140 (Oz) (range 88-176). Movement-related activity was analysed relative to force onset and referenced to AFz. The number of rowing strokes was on average (across subjects and conditions) 104 (range 84-117). The number of included trials (sites C3, C4, Cz, FCz) was on average 63 (range 20-117). Excluded trials were mainly contaminated by ocular artifacts. Due to the low number of channels an ICA-based artifact correction was not performed. The number of included trials in the movement-related condition was lower as in the VEP condition because of the longer segments (3 s vs. 1 s). One recording (subject D) was made on a hot August day. This resulted in large sweating artifacts, starting in condition three. Therefore, for the VEP analysis data of all recordings were filtered with a 1-Hz/24 dB highpass. This filtering did not visibly affect the morphology of the VEP waveform. For the analysis of the movement-related activity no additional highpass-filtering was made because at stroke rate 20/min movement frequency is about 0.33 Hz.

### 5.2 Results

Fig. 9a displays the VEPs generated by the standard stimulation (checkerboard) of the oddball task for all four subjects and conditions. For direct comparison, the VEP measured at active electrode O1 and passive electrode Oz are superimposed. As in the previous studies the VEPs were similar between rowing and rest, and the intraindividual differences were much smaller than the interindividual differences. Remarkably, the VEP measured at O1 and Oz were highly congruent. Fig. 9b displays the Standard Deviations (SD) of the EEG segments from which the average VEP was computed for all subjects and the six conditions. SDs were smallest in condition five (rest) and largest in the high intensive rowing condition four (except subject J). The signal-to-noise ratio (SNR) was computed in addition and revealed a similar pattern than the SD, that is the highest SNR in condition five and the lowest in condition four. However SD as well as SNR values could be misleading, if an average waveform is dominated by a large and systematic artifact. Nevertheless these data show that the qualitiy of the ERP is lower when movement intensity increases, as is indicated in the VEP waveforms as well.

Fig. 10 displays the motor-related activity at Cz. The waveforms show a systematic modulation by force output (and probably force-related artifacts) but not by movement speed (stroke rate). However the large interindividual differences are physiologically less plausible.

In summary, as shown in the previous studies, standard VEPs can be measured reliably during rowing whereas signal quality decreases when movement intensity increases. The differences between the VEP measured at active electrode O1 and passive electrode Oz are marginal to small. This shows that the use of active electrodes does not considerably suppress movement artifacts when electrode cables are shortened and fixated and a head-mounted amplifier is used. Consequently, the motor-related activity is still distorted by remaining artifacts.

## 6 General Discussion

The present pilot studies tested the practicability of four different EEG acquisition systems for ERP measures in moving subjects. Technically, including the modifications, the used systems were suited to measure ERPs in moving subjects in contrast to conventional laboratory EEG hardware with passive electrodes. The advantages of the used systems were the head-mounted amplifier (studies 1, 3, and 4) and the active electrodes (studies 2 and 4), both methods reducing cable movement artifacts. A further advantage of meanwhile available systems with low weight and small dimensions for movement research resp. research on moving subjects is either wireless data transmission or storing the data on a SD-card in the device itself.

The data from fourteen single case measures from six different subjects revealed for a standard paradigm (visual oddball task) comparable intraindividual ERPs during rowing and during rest (non-movement condition) in all cases, despite remaining artifacts in the data. EEG parameters and ERP waveforms are genetically determined and show generally a broad range of interindividual differences, but are also remarkably stable over time in adult subjects (Stassen et al., 1998; Weisbrod et al., 2004). This fact (although probably not well known), strongly supports the reliability of the data, because the intraindividual differences were much smaller than the interindividual differences. Higher intraindividual differences instead would indicate that the ERP pattern are largely distorted by movement artifacts.

Whereas the VEPs were quite similar, the P300 was smaller during rowing than during rest (studies 1-3). This may be partly due to a habituation effect (the rest condition was always performed before the rowing condition), like recently suggested by Scanlon et al. (2017) as well, who used also an oddball task to compare ERP measures during indoor cycling with a resting condition. Another contribution to this effect was probably that multiple task demands reduce the P300 (e.g., as reviewed by Polich, 2007). Here, attention is divided by counting the targets on the one hand and performing the rowing movement and the monitoring of stroke rate and power output on the other hand, which may be demanding for nonskilled rowers. Similar results with about a 30% smaller P300 amplitude during walking compared to sitting still (in counterbalanced order) were reported by Debener et al. (2012) and De Vos et al. (2014), who also suggested that the different task demands were the reason for this result. However, it has to be emphasised that the aim of the present pilot studies was not to investigate cognitive processes in rowing, instead these robust ERP components (VEPs, P300) were measured for methodological reasons, that is to compare ERP data quality during rowing and rest in a repeated measure design.

The positive results of the oddball task are promising for the investigation of brain functions in naturally behaving subjects outside the laboratory: for example in cognition research, brain-computer interface (BCI) applications, ambulatory assessment, and others. Since rowing is a very athletic sport and therefore a source of large movement-related artifacts, ERP measures with less motor activity such as walking around (e.g., Debener et al., 2012) or cycling when pedaling slowly at a subaerobic level (Scanlon et al., 2017) can easily be obtained using suitable equipment.

Due to the nature of ecological settings more limitations compared to laboratory settings have to be accepted. The placement of the stimulation monitor besides the ergometer and the oscillating viewing distance may be regarded as critical. Alternatives might be the use of a head-up display or an acoustical stimulation. However the gyroscope data in studies 3 and 4 revealed only small head movements in the oddball task during rowing, therefore it can be concluded that this was comparable in Study 1 and 2 because the placement of the monitor was the same. Eye movements were marginal or not present in the high densitiy recording of Study 2 but could be monitored at frontal sites in the other studies as well.

The (slight) differences in hardware and filter settings, and electrode positions were acceptable because reliability was assessed by the within-subject comparisons for each study separately.

The small sample size of these four pilot studies may be seen as critical from a classical cognitive neuroscience point of view. However it has to be noted that no subtle cognitive processes were investigated which require a larger sample size. Instead, robust electrophysiological (i) and physical (ii) processes were investigated which were seen in all fourteen measures. That is (i) VEPs were obtained during rowing and rest in all measures. And (ii) it was not possible to measure clean movement related neuronal activities. That is the chosen technical setup clearly differentiates between possibilities and limitations.

The second and more challenging aim of the four studies was to test if motor-related activity could be measured. This approach is unique for these pilot studies as the cited studies in the introduction aimed to measure cognitive processing in movement conditions, not the movement itself. Although motor potentials during the drive phase of the rowing movement, modulated by force output, were indicated (cf. Figure 6 and 10), in all four studies large movement-related artifacts occurred which distorted motor-related activity. These artifacts can be identified at electrode sites apart from the pre- and primary motor cortex. In this context it has to be considered that artifacts originating from the reference electrode will affect the other electrodes. As the classical mastoid reference captures EMG activity of head and neck muscles, reference electrode positions less affected by this EMG activity, as well as the activity of cortical motor areas, may be better suited (e.g. prefrontal sites or nose tip). At sites covering the motor areas artifacts are more difficult to detect because muscle force generation, movement kinematics, and movement-related artifacts have the same time course. Therefore, further technical improvements to reduce artifacts beforehand or to identify artifacts better (cf. Castermans et al., 2014) and correct them are necessary to investigate motor behavior in movements including the whole body (as in sports: for example, motor learning or differentiating high from low performance in movement execution). One example to identify artifact sources was given in Study 3 when using the gyroscope to identify artifacts generated by rapid head movements.

Known artifact sources are the EMG activity and sweating artifacts. The latter cannot be filtered out when movement frequency is in the same range (like in one subject of Study 4). Other sources of artifacts may rely on small movements of the electrode cables relative to the cap which were still possible; and the translational head movement during rowing in an electromagnetically noisy environment (the room was not shielded) may have generated small currents in the cables, as in a generator (according to Faraday’s law). However in both cases the active electrodes should be less vulnerable to this artifact sources. Furthermore, for the second case, artifacts should be larger when movement speed increases which was not observed.

To investigate further possible sources of motion artifacts, Oliveira et al. (2016) used a phantom head to simulate motion artifacts in EEG data and found that artifacts increased with movement frequency as well as with movement amplitude, that is in general with the acceleration of the phantom head. “We speculate that the major source of such artifacts is micro-movement of the recording electrodes in relation to the scalp surface” (Oliveira et al., 2016). Their data showed that artifacts strongly increased when the head acceleration was larger than 1.5 g. Based on these results, additional measures of head acceleration using a triaxial acceleration sensor (Move II, Movisens GmbH, Karlsruhe/Germany, www.movisens.com) attached to the Emotiv headset were analysed. These revealed values between 0.85 g in low intensive rowing (75 W, 20 strokes/min) and 2.5 g in high intensive rowing (360 W, 30 strokes/min). That is the lower quality of VEPs in Study 4 at the rowing conditions with higher intensity may partly depend on such micro-movements of the electrodes, independently if passive or active electrodes are used because the main advantage of active electrodes is that these are less susceptible for cable sway artifacts.

Another pitfall with a physiological origin might be, that different neuronal activities are superimposed which hinders the detection of relevant activities. Siemonow et al. (2000) observed that the amplitude of the motor potential correlates very high (r = 0.93) with force output in an isometric elbow flexion task and reported values of up to 8 μV/150 N. In rowing these values (assuming a physiological source) are considerably higher (about 100 μV and 1000 N in Study 4). In contrast to these, probably from the pyramidal cells in the primary motor cortex generated efferent activities (Brecht et al., 2004; Shibasaki, 2012), are other motor-related activities (originating from the premotor cortex or the SMA) very small. E.g. in two own visuomotor tracking studies the effects related to motor learning were below 1 μV (Hill, 2009, 2014). That is, if generally the activities related to motor learning or motor programming (e.g. in rowing the perception and adaptation of within-crew differences of rowing technique, Hill, 2002) are in this amplitude range, these will be hidden when the activity related to force execution is much higher.

## References

Atsumori, H., Kiguchi, M., Katura, T., Funane, T., Obata, A., Sato, H., et al. (2010). Noninvasive imaging of prefrontal activation during attention-demanding tasks performed while walking using a wearable optical topography system. Journal of Biomedical Optics, 15, 046002-1-7.

Barham, M.P., Clark, G.M., Hayden, M.J., Enticott, P.G., Conduit, R., Lum, J.A.G. (2017). Acquiring research-grade ERPs on a shoestring budget: A comparison of a modified Emotiv and commercial SynAmps EEG system. Psychophysiology, 54(9), 1393–1404. doi: 10.1111/psyp.12888.

Badcock, N.A., Mousikou, P., Mahajan, Y., de Lissa, P., Thie, J., McArthur, G. (2013). Validation of the Emotiv EPOC(®) EEG gaming system for measuring research quality auditory ERPs. PeerJ., 1:e38. doi: 10.7717/peerj.38

Bertrand, O., Perrin, F., Pernier, J.A. (1985). A theoretical justification of the average reference in topographic evoked potential studies. Electroencephal. Clin. Neurophysiol., 62, 462–464.

Brecht, M., Schneider, M., Sakmann, B., Margrie, T. W. (2004). Whisker movements evoked by stimulation of single pyramidal cells in rat motor cortex. Nature, 427, 704–710.

Castermans, T., Duvinage, M., Cheron, G., Dutoit, T. (2014). About the cortical origin of the low-delta and high-gamma rhythms observed in EEG signals during treadmill walking. Neurosci. Lett., 21, 166–170. doi: 10.1016/j.neulet.2013.12.059.

Chi, Y.M., Wang, Y.T., Wang, Y., Maier, C., Jung, T.P., Cauwenberghs, G. (2012). Dry and noncontact EEG sensors for mobile brain-computer interfaces. IEEE Trans. Neural. Syst. Rehabil. Eng., 20, 228–35. doi: 10.1109/TNSRE.2011.2174652.

Dai, T.H., Liu, J.Z., Sahgal, V., Brown, R.W., Yue, G.H. (2001). Relationship between muscle output and functional MRI-measured brain activation. Exp. Brain. Res., 140, 290–300.

Debener, S., Minow, F., Emkes, R., Gandras, K., de Vos, M. (2012). How about taking a low-cost, small, and wireless EEG for a walk? Psychophysiology, 49, 1449–53.

De Vos, M., Gandras, K., Debener, S. (2014). Towards a truly mobile auditory brain-computer interface: Exploring the P300 to take away. Int. J. Psychophysiol., 91, 46–53. doi: 10.1016/j.ijpsycho.2013.08.010.

Dum, R.P., and Strick, P.L. (2002). Motor areas in the frontal lobe of the primate. Physiology & Behavior, 77, 677–682.

Duvinage, M., Castermans, T., Petieau, M., Hoellinger, T., Cheron, G., Dutoit, T. (2013). Performance of the Emotiv Epoc headset for P300-based applications. Biomed. Eng. Online, 12, 56. doi: 10.1186/1475-925X-12-56.

Enders H, Nigg BM (2016). Measuring human locomotor control using EMG and EEG: Current knowledge, limitations and future considerations. Eur J Sport Sci., 16(4), 416–26. doi: 10.1080/17461391.2015.1068869. Epub 2015 Aug 4.

Enders H, Cortese F, Maurer C, Baltich J, Protzner AB, Nigg BM (2016). Changes in cortical activity measured with EEG during a high-intensity cycling exercise. Journal of Neurophysiology, 115, 379–338.

Ferree, T.C., Luu, P., Russell, G.S., Tucker, D.M. (2001). Scalp electrode impedance, infection risk, and EEG data quality. Clin. Neurophysiol., 112, 536–544.

Gevins, A., Chan, C.S., Sam-Vargas, L. (2012). Towards measuring brain function on groups of people in the real world. Plos One, 7, 1–9. doi: 10.1371/journal.pone.0044676.

Gramann, K., Gwin, J.T., Bigdely-Shamlo, N., Ferris, D.P., Makeig, S. (2010). Visual evoked responses during standing and walking. Front. Hum. Neurosci., 4, 202

Gramann, K., Gwin, J.T., Ferris, D.P., Oie, K., Jung, T.P., Lin, C.T., et al. (2011). Cognition in action: imaging brain/body dynamics in mobile humans. Rev. Neurosci., 22, 593–608. doi: 10.1515/RNS.2011.047.

Gramann, K., Ferris, D.P., Gwin, J., Makeig, S. (2014). Imaging natural cognition in action. Int. J. Psychophysiol., 91, 22–9. doi: 10.1016/j.ijpsycho.2013.09.003.

Gwin, J.T., Gramann, K., Makeig, S., Ferris, D.P. (2010). Removal of movement artifact from high-density EEG recorded during walking and running. J. Neurophysiol., 103, 3526–34.

Gwin, J.T., Gramann, K., Makeig, S., Ferris, D.P. (2011). Electrocortical activity is coupled to gait cycle phase during treadmill walking. Neuroimage, 54, 1289–96.

Hazeltine, E., and Ivry, R.B. (2002). Can We Teach the Cerebellum New Tricks? Science, 296, 1979–1980.

Hill, H. (2002). Dynamics of coordination within elite rowing crews: evidence from force pattern analysis. J. Sports. Sci., 20, 101–117.

Hill, H. (2009). An event-related potential evoked by movement planning is modulated by performance and learning in visuomotor control. Exp. Brain. Res., 195, 519–529.

Hill, H. (2014). Modulation of frontal and parietal neuronal activity by visuomotor learning. An ERP analysis of implicit and explicit pursuit tracking tasks. Int. J. Psychophysiol., 91, 212–24. doi: 10.1016/j.ijpsycho.2013.12.007

Kranczioch, C., Zich, C., Schierholz, I., Sterr, A. (2014). Mobile EEG and its potential to promote the theory and application of imagery-based motor rehabilitation. Int. J. Psychophysiol., 91, 10–5. doi: 10.1016/j.ijpsycho.2013.10.004.

Makeig, S., Bell, A.J., Jung, T.P., Sejnowski, T.J. (1996). Independent component analysis of electroencephalographic data. Adv. Neural. Inf. Process. Syst., 8, 145–151.

Oliveira, A.S., Schlink, B.R., Hairston, W.D., König, P., Ferris, D.P. (2016). Induction and separation of motion artifacts in EEG data using a mobile phantom head device. J. Neural. Eng., 13, 036014. DOI: 10.1088/1741-2560/13/3/036014

Picton, T.W., Bentin, S., Berg, P., Donchin, E., Hillyard, S.A., Johnson, J.R., et al. (2000). Guidelines for using human event-related potentials to study cognition: recording standards and publication criteria. Psychophysiology, 37, 127–152.

Piper, S.K., Krueger, A., Koch, S.P., Mehnert, J., Habermehl, C., Steinbrink, J., et al. (2014). A wearable multi-channel fNIRS system for brain imaging in freely moving subjects. Neuroimage, 85, 64–71. DOI: 10.1016/j.neuroimage.2013.06.062

Polich, J. (2007). Updating P300: an integrative theory of P3a and P3b. Clin. Neurophysiol., 118, 2128–2148.

Ranganathan, V.K., Siemionov, V., Liu, J.Z., Sahgal, V., Yue, G.H. (2004). From mental power to muscle power - gaining strength by using the mind. Neuropsychologia, 42, 944–956.

Ruffini, G., Dunne, S., Farres, E., Cester, I., Watts, P.C., Silva, S.P., et al. (2007). ENOBIO dry electrophysiology electrode; first human trial plus wireless electrode system. Conf. Proc. IEEE Eng. Med. Biol. Soc. 6690–4.

Scanlon, J.E.M., Sieben, A.J., Holyk, K.R., Mathewson, K.E. (2017). Your brain on bikes: P3, MMN/N2b, and baseline noise while pedaling a stationary bike. Psychophysiology, 54, 927–937. DOI: 10.1111/psyp.12850

Shibasaki H (2012). Cortical activities associated with voluntary movements and involuntary movements. Clinical Neurophysiology, 123, 229–243.

Siemionow, V., Yue, G.H., Ranganathan, V.K., Liu, J.Z., Sahgal, V. (2000). Relationship between motor activity-related cortical potential and voluntary muscle activation. Exp. Brain. Res., 133, 303–311.

Simon, O. (1977). Das Elektroenzephalogramm. München Wien Baltimore: Urban und Schwarzenbeck.

Stassen, H.H., Bomben, G., Hell, D. (1998). Familial brain wave patterns: study of a 12-sib family. Psychiatr. Genet., 8, 141–53.

Stone, D. B., Tamburro, G., Fiedler, P., Haueisen, H., Comani, S. (2018). Automatic Removal of Physiological Artifacts in EEG: The Optimized Fingerprint Method for Sports Science Applications. Front. Hum. Neurosci., 12, https://doi.org/10.3389/fnhum.2018.00096

Stopczynski, A., Stahlhut, C., Petersen, M.K., Larsen, J.E., Jensen, C.F., Ivanova, M.G., et al. (2014). Smartphones as pocketable labs: Visions for brain imaging and neurofeedback. Int. J. Psychophysiol., 91, 54–66. http://dx.doi.org/10.1016/j.ipsycho.2013.08.007

Weisbrod, M., Hill, H., Sauer, H., Niethammer, R., Guggenbühl, S., Hell, D., Stassen, H.H. (2004). Nongenetic Pathologic Developments of Brain Wave Patterns in Monozygotic Twins Discordant and Concordant for schizophrenia. Am. J. Med. Genet. B Neuropsychiatr. Genet., 125B, 1–9.

Wulf, G., and Shea, C.H. (2002). Principles derived from the study of simple skills do not generalize to complex skill learning. Psychomotoric Bulletin & Review, 9, 185–211.

Zschocke, S. (2002). Klinische Elektroenzephalographie. Berlin Heidelberg New York: Springer.

